# Exploring group-specific technical variation patterns of single-cell data

**DOI:** 10.1101/2024.09.20.614043

**Authors:** Yang Zhou, Qiongyu Sheng, Shuilin Jin

**Author notes:** Correspondence to Shuilin Jin.

## Abstract

Constructing single-cell atlases requires preserving differences attributable to biological variables, such as cell types, tissue origins, and disease states, while eliminating batch effects. However, existing methods are inadequate in explicitly modeling these biological variables. Here, we introduce SIGNAL, a general framework designed to disentangle biological and technical effects by learning group-specific technical variation patterns, thereby linking these metadata to data integration. SIGNAL employs a novel variant of principal component analysis (PCA) to align multiple batches, enabling the integration of 1 million cells in approximately 2 minutes. SIGNAL, despite its computational simplicity, surpasses state-of-the-art methods across multiple integration scenarios: (1) heterogeneous datasets, (2) cross-species datasets, (3) simulated datasets, (4) integration on low-quality cell annotations, and (5) reference-based integration. Furthermore, we demonstrate that SIGNAL accurately transfers knowledge from reference to query datasets. Notably, we propose a self-adjustment strategy to restore annotated cell labels potentially distorted during integration. Finally, we apply SIGNAL to multiple large-scale atlases, including a human heart cell atlas containing 2.7 million cells, identifying tissue- and developmental stage-specific subtypes, as well as condition-specific cell states. This underscores SIGNAL’s exceptional capability in multi-scale analysis.

## Introduction

The rapid evolution of single-cell technologies has empowered us to delve into extensive datasets, providing unprecedented insights into cell subtypes and functional intricacies. Large-scale single-cell atlases from various developmental stages, disease conditions, and tissues have been collected, integrated, and annotated (1-3). Constructing such atlases necessitates integration methods capable of preserving differences across diverse biological variables (hereafter referred to as ‘group variables’), beyond just cell types. However, this remains challenging due to the inherent difficulty in distinguishing meaningful biological variations from batch effects across different conditions.

A recent benchmark study (4) has demonstrated that integration methods incorporating cell type labels outperform fully unsupervised approaches, highlighting the critical role of metadata in data integration. Current methods such as Harmony (5) and scVI (6) capitalize on other meta information, such as tissue or developmental stage labels, to facilitate integration. However, these methods treat such metadata as ancillary batch variables, seeking to mitigate rather than explicitly preserve the introduced differences. For example, Suo et al. (3) employed scVI, utilizing the library construction protocols (5’ and 3’) and donor labels as batch variables to retain the differences between organs and developmental stages. In contrast, other popular unsupervised methods (7-12) typically disregard these biological variables, reducing all inter-batch differences to batch effects, a strategy prone to over-correction.

Here, we present SIGNAL (SIngle-cell Group techNical vAriations Learning), a computationally simple yet effective framework that leverages metadata to guide data integration. In addition to cell type labels, SIGNAL can utilize other group variables to integrate data in a manner that preserves differences among subgroups defined by these variables. By group technical variations (GTVs), SIGNAL can transfer knowledge from reference to query datasets, enabling cell label prediction and reference-based integration tasks. We apply SIGNAL to various integration scenarios across both simulated and real datasets, demonstrating its superior integration performance compared to state-of-the-art methods. Furthermore, we demonstrate that SIGNAL is particularly well-suited for the construction and multi-scale analyses of customized single-cell atlases.

## Results

### Overview of SIGNAL

SIGNAL aims to align subgroups across different batches while preserving differences between them, guided by a given group variable such as cell type or condition (Fig. 1A). When the group variable is cell type, SIGNAL operates as a supervised integration method. For more complex analytical needs, SIGNAL can incorporate multiple group variables (Fig. 1A and *Materials and Methods*).

**Figure 1.**
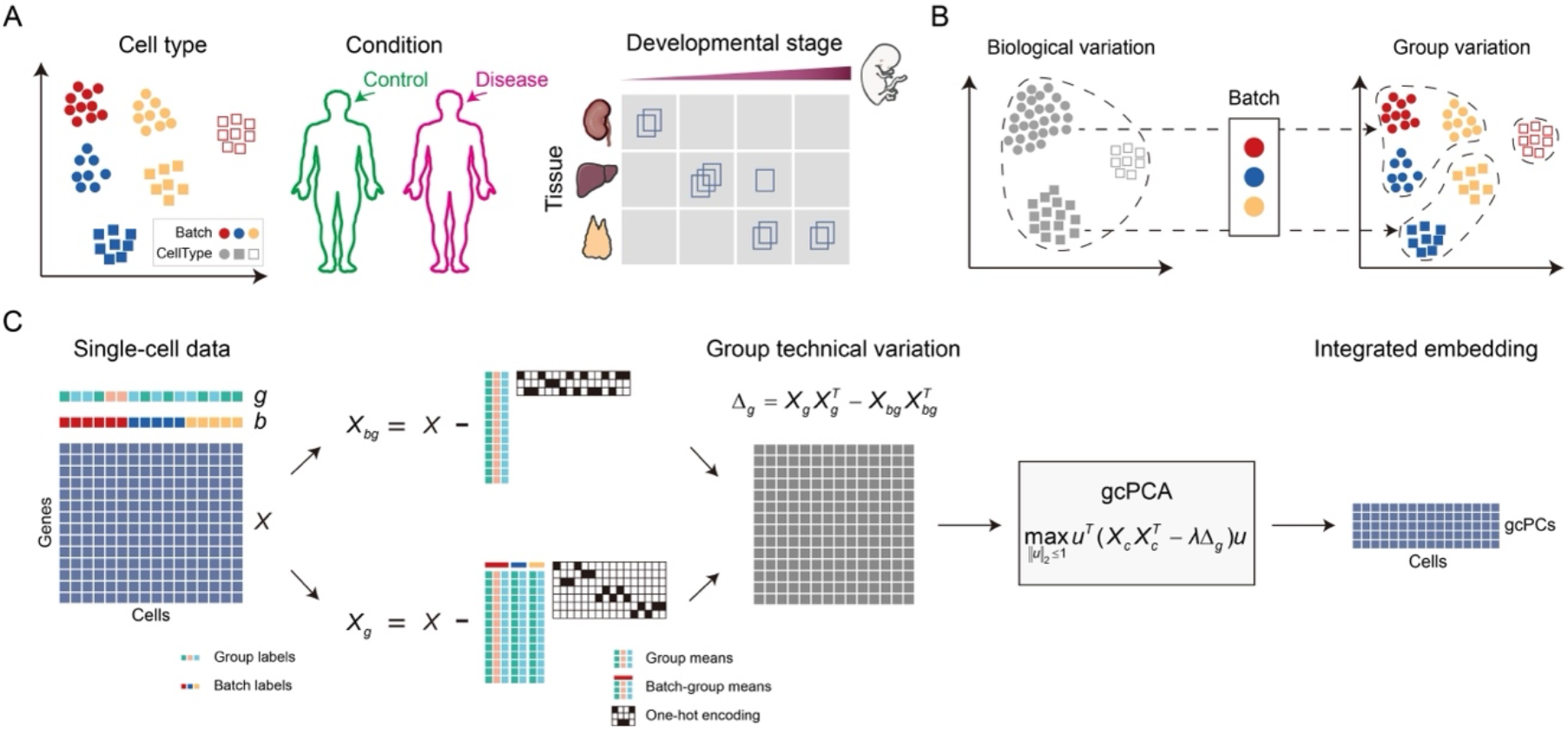
SIGNAL framework overview. (A) Examples of group variables. Single-group settings (left and middle) and two-group settings (right) are available in the SIGNAL framework. (B) Estimating technical variation using group variation and group biological variation. (C) SIGNAL integration. SIGNAL learns GTVs and projects the data into the gcPC space using gcPCA.

The rationale behind SIGNAL lies in decomposing total variation into biological and technical variations, similar to our previous work on single-cell RNA sequencing (scRNA-seq) data integration (13). Batch effects are considered to act on each individual subgroup, and these subgroup-specific technical variations can be calculated through variation decomposition (Fig. 1B and Fig. S1). Therefore, we propose using GTVs to estimate the technical variation in the dataset. Given a single-cell data matrix *X* along with one or multiple group variables derived from the metadata, SIGNAL learns GTVs, which serve as inputs for downstream tasks including data integration and reference-based label prediction (Fig. 1C and Fig. S2). Specifically, GTV is the difference between 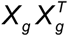 and 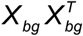, referred to as group variation and group biological variation, respectively, representing the total technical variation of all subgroups introduced by batch variables.

To integrate data using GTVs, we introduce group-centered principal component analysis (gcPCA), a variant of classical PCA by subtracting the GTVs from the total variation and learning a set of projection vectors to project data onto a common low-dimension space (Fig. 1C and *Materials and Methods*). For transferring knowledge from an annotated dataset to a newly arrived unannotated dataset, we developed a reference-based cell label prediction method called group label prediction using group technical variation (PGTV) (Fig. S2A). After assigning cell labels from the reference dataset to the query dataset, these datasets can be aligned using gcPCA, with an updated variation matrix (Fig. S2B).

### Benchmark of SIGNAL and state-of-the-art integration methods

To evaluate the integration performance of SIGNAL, we benchmarked it against state-of-the-art methods. We curated nine single-cell datasets, including eight datasets collected for benchmarking integration methods (4, 14) and one from a recent cross-species study (15). These datasets were organized into three integration tasks: heterogeneous datasets, cross-species datasets, and simulated datasets, representing a broad range of data integration scenarios. In addition to SIGNAL, nine competitors were employed, encompassing the most representative data integration methods. These ranged from statistical methods based on similar cells or shared cell types (fastMNN (7), Seurat (8), Harmony, and LIGER (11)) to graph-based methods (BBKNN (16) and Conos (17)) and deep learning methods (scVI, scANVI (18), and scPoli (19)). Ground-truth cell type labels were provided to scANVI, scPoli, and SIGNAL to guide integration. We assessed the integration results using metrics proposed by Luecken et al. (4), which evaluate biological variation conservation and batch effect removal, and computed overall integration scores (*Materials and Methods*).

We found that SIGNAL and scPoli excelled in both biological variation conservation and batch effect removal (Fig. 2A), with SIGNAL achieving the highest average scores. In overall integration performance, the three methods utilizing cell type labels ranked highest (Fig. 2B), aligning with the observations made by Luecken et al. (4). Specifically, SIGNAL ranked third on the immune (cross-species) dataset and within the top two on the remaining eight datasets, demonstrating consistently superior integration performance. Visualization using uniform manifold approximation and projection (UMAP) (20) showed that SIGNAL well mixed cells from multiple batches and distinguished different cell types across all datasets (Figs. S3-S5). In contrast, scANVI failed to align cell types correctly in the simulated datasets, while scPoli struggled with the human dendritic cells (DCs) dataset (with similar cell subtypes across batches) and the white adipose tissue (WAT) dataset (multi-batch and cross-species).

**Figure 2.**
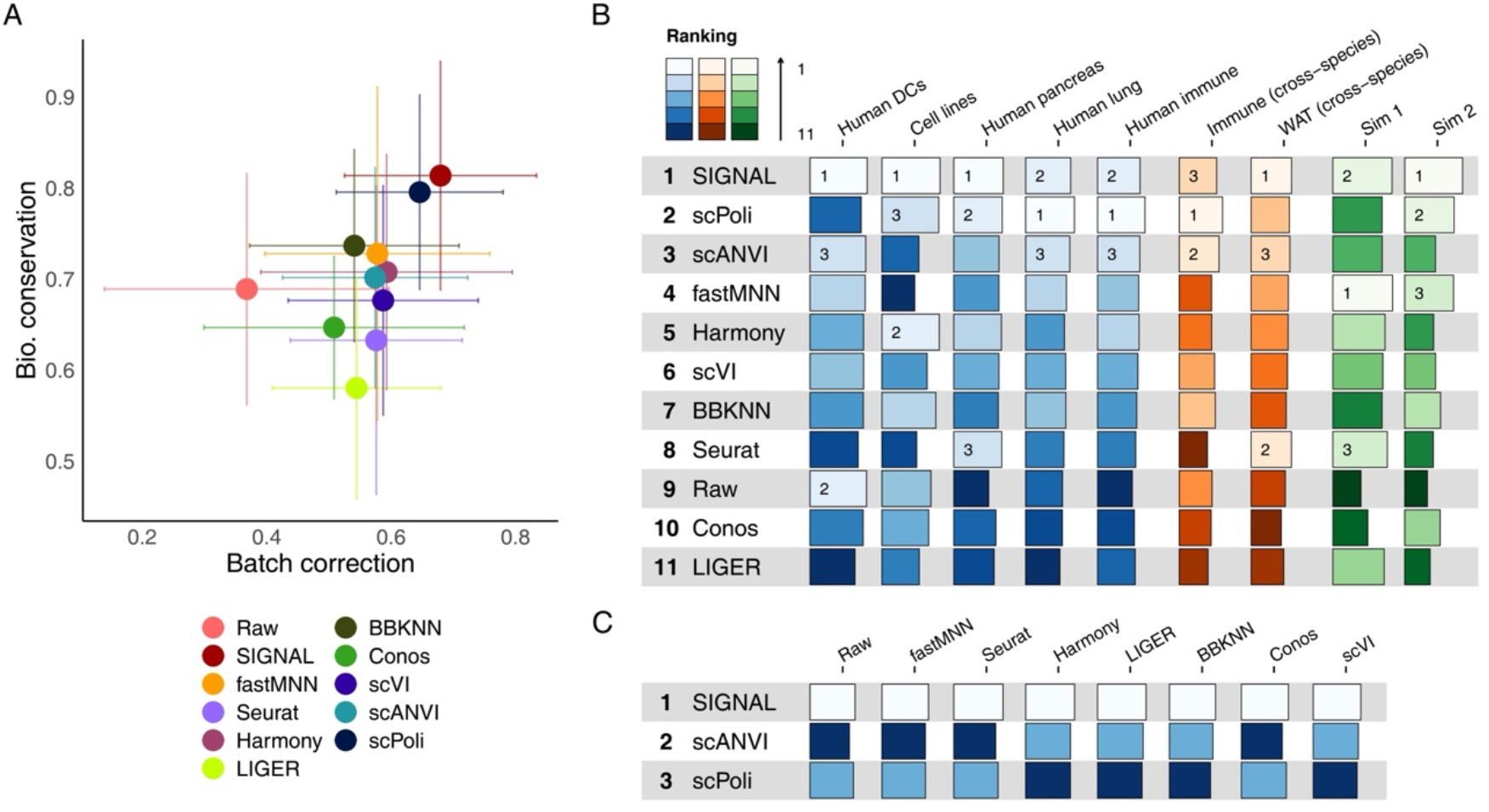
SIGNAL outperforms state-of-the-art integration methods. (A) Scatter plot of biological variation conservation and batch correction performance of various integration methods across nine scRNA-seq datasets. Error bars indicate the standard deviation across datasets. (B) Overall ranking of integration scores for the nine datasets. The top three methods for each dataset are indicated. (C) Overall ranking of integration scores for the integration tasks involving low-quality labels.

Annotating single-cell datasets is often time-consuming and labor-intensive. To be broadly applicable, supervised data integration methods should perform well even with low-quality labels. To test this, we used the human pancreas dataset, containing multiple batches from different sequencing technologies (*Materials and Methods*). We subjected the integrated results by various unsupervised methods to k-means clustering to simulate a rough cell label annotation process. The resulting annotations were then used as inputs for scPoli, scANVI, and SIGNAL integration. Despite the low-quality labels, all three methods yielded satisfactory integration results (Fig. 2C and Fig. S6). Notably, SIGNAL consistently outperformed the other methods, demonstrating remarkable robustness to input label quality.

### SIGNAL predicts cell labels based on reference annotations

We next investigated the transfer of knowledge from an annotated reference dataset to a query dataset. We evaluated the label prediction performance of PGTV against common reference-based annotation methods, including SingleR (21), SciBet (22), scmap (23), and scPred (24), across two simulated and four real datasets (*Materials and Methods*). PGTV demonstrated label prediction performance comparable to SingleR and SciBet in terms of accuracy and macro F1 scores, outperforming other methods across all datasets (Fig. S7).

PGTV relies on reference cell type annotations. Considering the low-quality labels, we introduced a self-adjustment strategy for PGTV, which iteratively predicts and adjusts cell labels within the dataset (*Materials and Methods*). We simulated a mislabeled dataset by manually altering cell labels in the Sim 1 dataset (Fig. 3A). The PGTV-adjusted results for each individual batch and across all batches showed that PGTV accurately recovered the ground truth cell labels (Fig. 3B).

**Figure 3.**
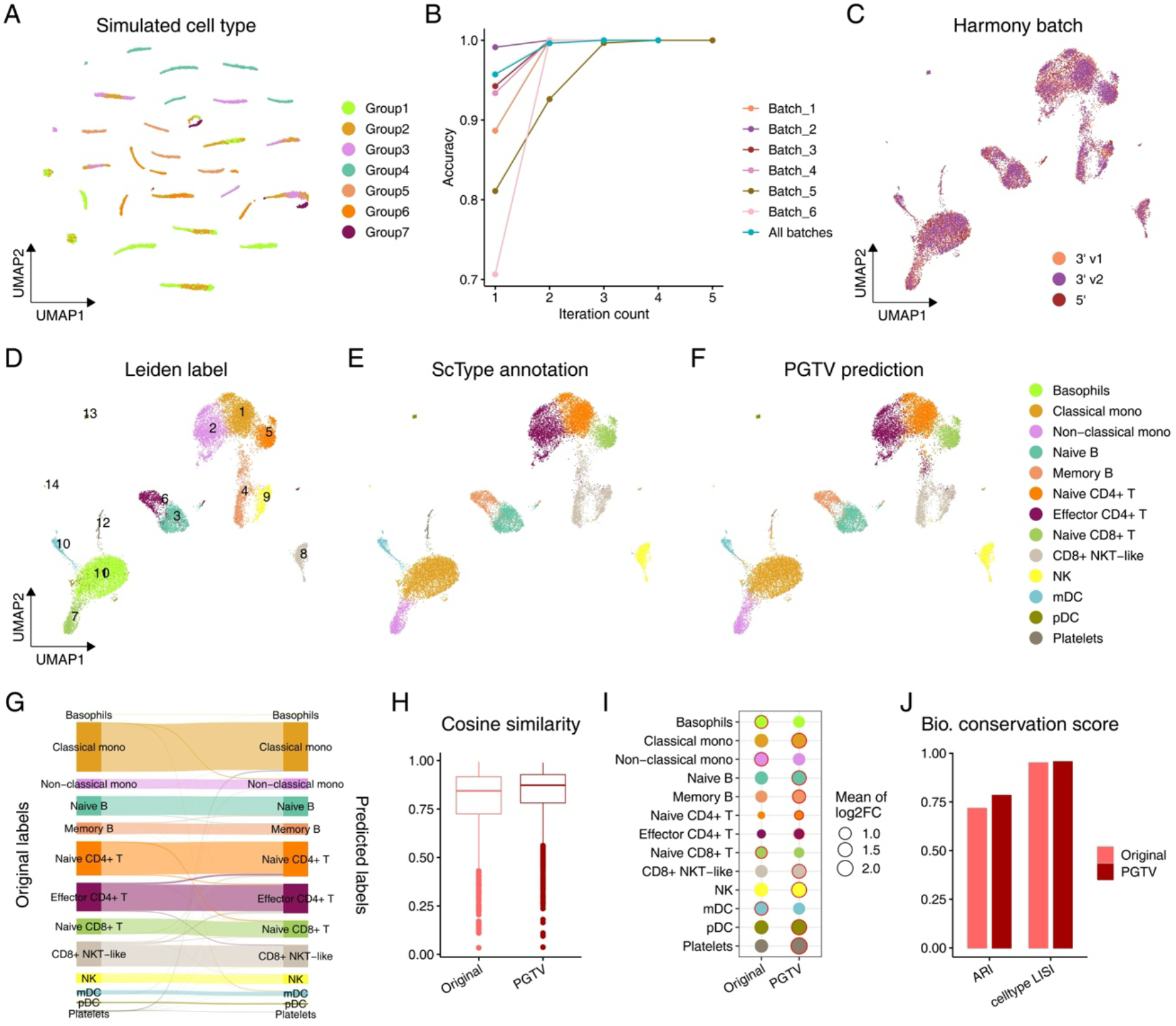
PGTV adjusts cell labels. (A) UMAP visualization of the Sim 1 dataset, with cells colored by simulated mislabeled cell type. (B) Accuracy of PGTV-adjusted results across different numbers of iterations. (C-F) UMAP visualization of the Harmony-integrated human PBMCs (multi-platform) dataset, with cells colored by batch (C), Leiden label (D), ScType-annotated label (E), and PGTV-adjusted label (F). (G) PGTV-adjusted result. (H) Cosine similarities between cells with inconsistent labels and cells with consistent labels. (I) Comparison of the mean log-fold change of the top 5 markers for cell types between original and PGTV-predicted labels. Results with a larger mean log-fold change of the same cell type are highlighted with red circles. (J) Biological conservation scores of SIGNAL-integrated results using original and PGTV-adjusted labels.

We further applied PGTV self-adjustment to a human peripheral blood mononuclear cells (PBMCs) dataset, containing three batches with a total of 20,517 cells. We annotated the dataset using a combined strategy involving Harmony integration, Leiden clustering (25), and ScType annotation (26), identifying 13 cell types (Fig. 3 C-E). This common pipeline in integrative analysis, though effective, can lead to misannotated cells due to its dependence on the integration method. PGTV reassigned a portion of cells into new labels, primarily within similar cell populations such as T cell subtypes (Fig. 3 F and G). We compared the cosine similarities between cells with discrepant annotations and their different assigned labels (*Materials and Methods*). PGTV-adjusted labels exhibited higher similarities (Fig. 3H). Additionally, we examined the differential expression on original and PGTV-adjusted labels. Marker genes for cell types showed higher log-fold changes in PGTV-adjusted labels (Fig. 3I), indicating that PGTV effectively recovered the biological heterogeneity of cell types. As demonstrated previously, SIGNAL is robust to low-quality cell labels, enabling the evaluation of annotation reliability using SIGNAL-integrated results. We compared the biological conservation scores of original and PGTV-adjusted labels and found that integrated results on PGTV-adjusted labels achieved higher ARI and cell type LISI (local inverse Simpson’s index) values than original labels (Fig. 3J). These results together showed that PGTV can effectively restore cell labels distorted by data integration.

### SIGNAL enables reference-base integration

We further assessed SIGNAL’s capability of reference-based integration in comparison to state-of-the-art methods, including scANVI, scPoli, Seurat v4 (27), and Symphony (28) (Fig. 4). Effective reference-based integration should seamlessly merge query data with reference data while accurately transferring cell labels. We evaluated dataset mixing using dataset LISI and assessed label transfer performance through accuracy and macro F1 scores (*Materials and Methods*).

**Figure 4.**
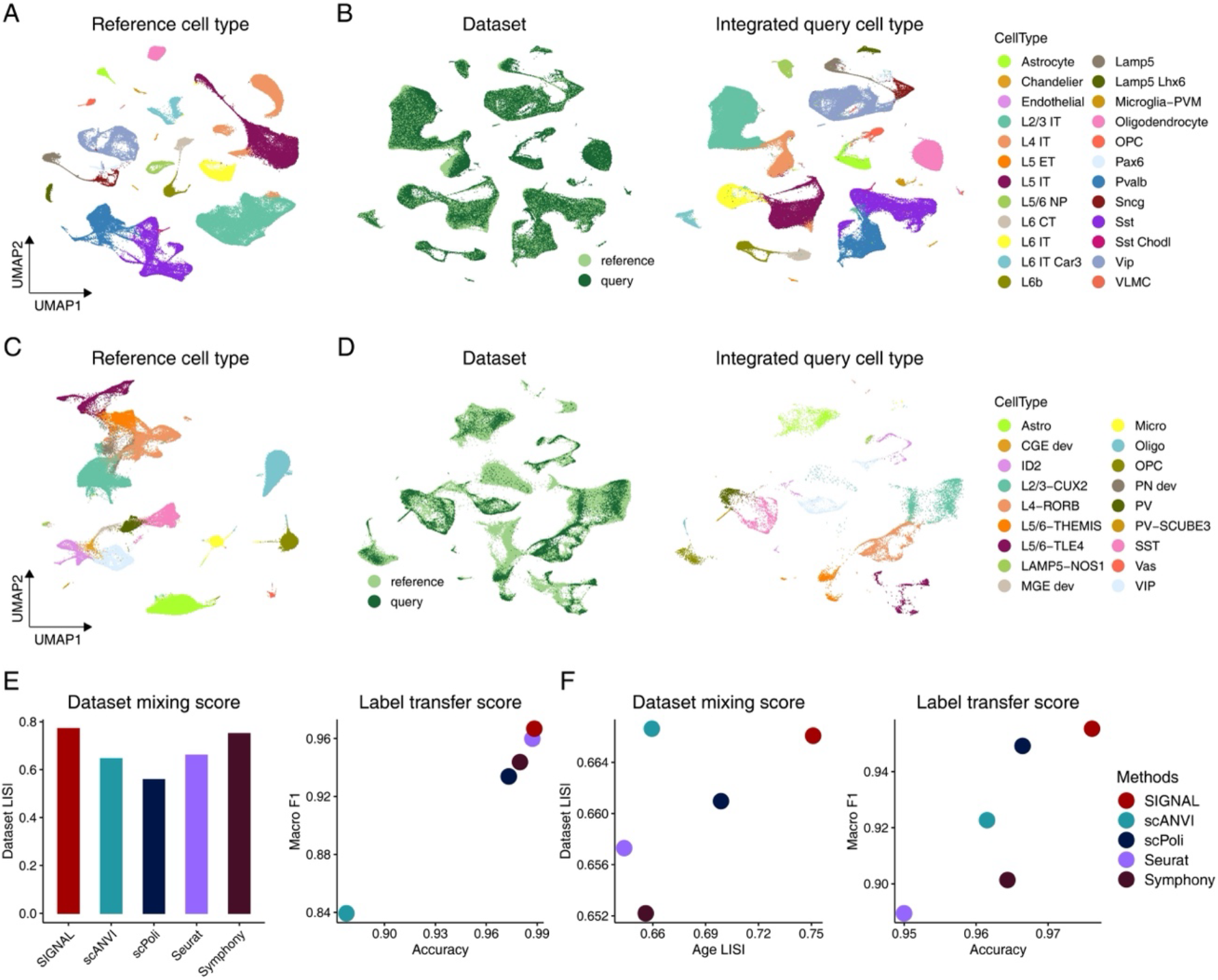
SIGNAL integrates query with reference dataset. (A) UMAP visualization of the integrated human MTG dataset, colored by cell type. (B) UMAP visualizations of the integrated results for the reference human MTG dataset and query human cortex dataset, colored by dataset (left) and query cell type (right). (C) UMAP visualization of the integrated reference human cortical development dataset, colored by cell type. (D) UMAP visualizations of the integrated results for the reference and query dataset of the human cortical development data, colored by dataset (left) and query cell type (right). (E), (F) Comparison of dataset mixing metrics and label transfer metrics for the human cortex (E) and human cortical development (F) datasets.

First, we tested whether SIGNAL could effectively integrate multi-batch reference and query datasets from different studies. In this task, the human middle temporal gyrus (MTG) dataset (29) (containing five batches and 137,303 nuclei) served as the reference, and the human cortical dataset (30) (containing 78 batches and 379,330 nuclei) was the query, exhibiting strong batch effects between datasets (Fig. S8A). SIGNAL integration distinctly separated different cell types within the reference dataset (Fig. 4A). For the query dataset, PGTV achieved a cell type prediction accuracy of 0.97 and a Macro F1 score of 0.96 (Fig. S8B). SIGNAL then used the predicted labels to integrate the reference and query datasets, successfully mixing them while clearly distinguishing cell types within the query dataset (Fig. 4B). In contrast, scANVI, scPoli, and Seurat struggled to mix the datasets effectively, while Symphony produced a joint embedding that mingled distinct cell types (Fig. S8 C-F). Comparatively, SIGNAL achieved the best performance in both dataset mixing and label transfer (Fig. 4E).

A crucial application of reference-based integration is handling developmental data, which presents the challenge of maintaining developmental structures. We applied SIGNAL to a human cortical developmental dataset (31), divided into reference (containing 24 batches and 140,117 nuclei) and query (containing three batches and 13,356 nuclei) based on 10x 3’ v3 and v2 chemistry, respectively. SIGNAL succeeded in integrating the reference data, preserving all cell identities (Fig. 4C and Fig. S9A). Based on predicted high-precision query cell labels (Fig. S9B), SIGNAL integration well mixed the reference and query datasets while maintaining developmental stage differences in broad cell types (Fig. 4D and Fig. S9C). Notably, for L5/6-THEMIS and L5/6-TLE4 subtypes in the query, SIGNAL accurately positioned them within the postnatal reference cells, distinguishing them from prenatal cells, indicating significant expression differences before and after birth. In contrast, other methods mixed the reference and query datasets effectively but failed to maintain developmental stage differences (Fig. S9 D-G). For example, L5/6-THEMIS and L5/6-TLE4 cells from different stages were intermingled. To evaluate the ability of integration methods to preserve stage differences, we introduced age LISI, which reflects the degree of separation of cells from different stages in the integrated results (*Materials and Methods*). We found that SIGNAL outperformed other methods in both dataset mixing and label transfer, leading in overall performance (Fig. 4F).

In reference-based integration, a common scenario is the presence of novel cell types in the query dataset. To this end, we computed confidence scores for cell label predictions and excluded query cells with low confidence scores during the reference-based integration (*Materials and Methods*). To evaluate the effectiveness of this approach, for the reference-based integration task on human cortical data, we removed the cell types with the lowest and highest abundances in the query from the reference dataset. All query-specific cell types exhibited low confidence scores and could be distinctly identified in the SIGNAL-integrated results (Fig. S10). This emphasizes the power of SIGNAL to discover and accurately integrate novel cell types in query data.

### SIGNAL enables multi-scale analysis of large-scale single-cell atlas

We next demonstrated how SIGNAL can be applied to the construction and multi-scale analyses of customized single-cell atlases, addressing increasingly complex application scenarios. First, we applied SIGNAL to the recently published human lung cell atlas (HLCA) (32). Using the well-constructed HLCA core (584,944 cells from 14 datasets) as the reference, we sought to transfer its high-resolution cell type annotations to the multi-source HLCA extension (1,074,190 cells from 36 datasets). The query cells exhibited high confidence scores and were well integrated with the reference cells (Fig. 5 A and B and Fig. S11A). While scANVI and scPoli also achieved good integration of reference and query cells (Fig. S11 B and C), SIGNAL outperformed these methods in label transfer performance, achieving an accuracy of 0.87 and a macro F1 score of 0.73 (Fig. 5C). This highlights SIGNAL’s superior performance in transferring cell annotations in complex integration tasks.

**Figure 5.**
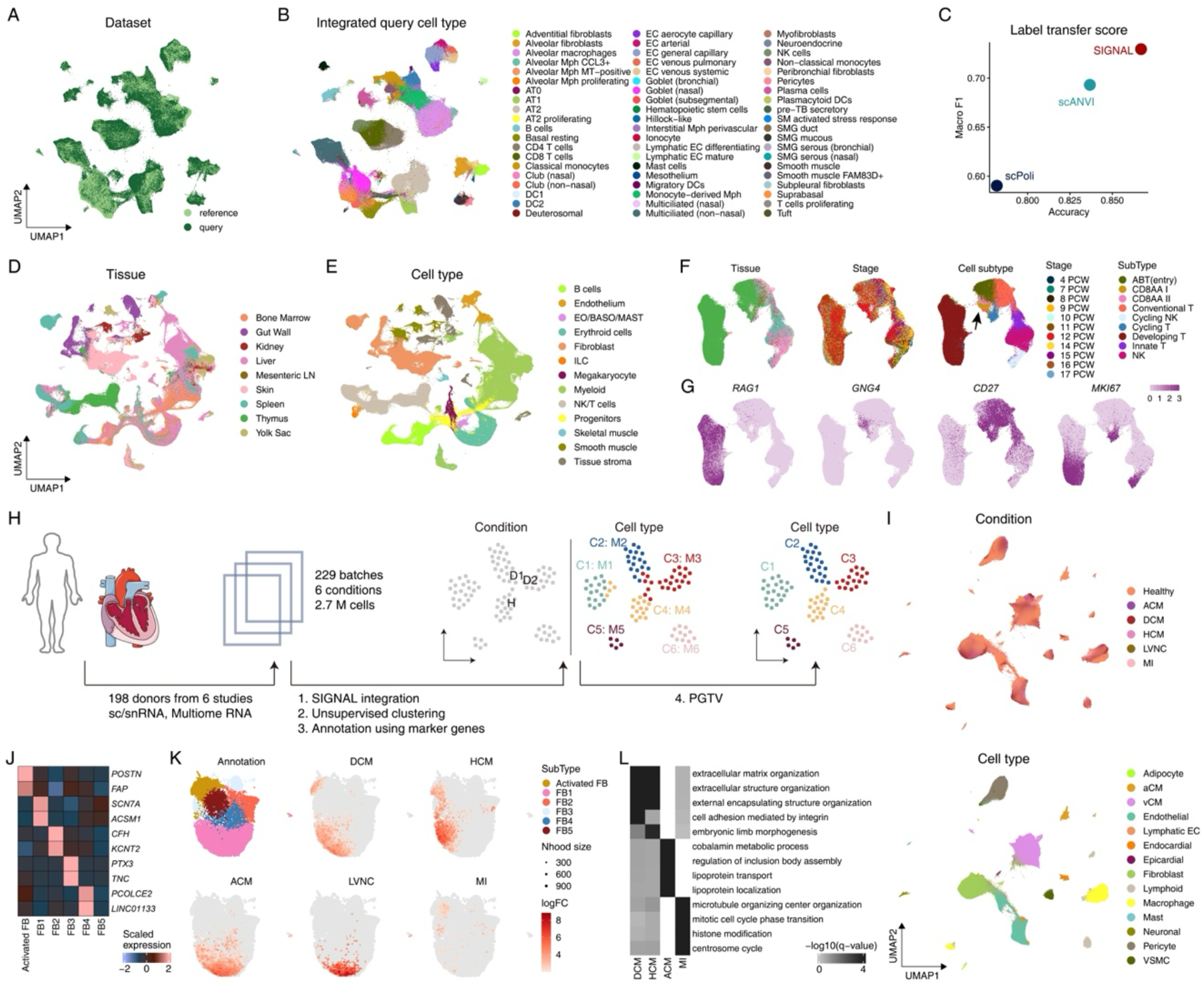
SIGNAL enables multi-scale analysis of large-scale single-cell atlas. (A), (B) UMAP visualizations of the integrated results for the reference (HLCA core) and query (HLCA extension), with cells colored by dataset (A) and query cell type (B). (C) Comparison of label transfer metrics for the integrated HLCA. (D), (E) UMAP visualizations of the SIGNAL-integrated developing human immune cell atlas, with cells colored by tissue (D) and cell type (E). (F) UMAP visualizations of NK/T cells in SIGNAL-integrated data, with cells colored by tissue (left), stage (middle), and cell subtype (right). (G) Expression patterns of marker genes for NK/T cell subtypes. (H) Overview of human heart cell atlas construction. (I) UMAP visualizations of SIGNAL-integrated human heart cell atlas, with cells colored by condition (top) and cell type (bottom). (J) Expression patterns of marker genes for heart fibroblast subtypes. (K) Milo differential abundance results. Each point represents a neighborhood, colored by fibroblast subtype and log-fold change in abundance between the specified condition and healthy. (L) Overrepresented biological processes of activated fibroblasts across conditions.

Next, using a multi-tissue, multi-developmental stage developing human immune cell atlas (3), we tested whether SIGNAL could identify tissue-and stage-specific cell subtypes. By treating tissue and developmental stage as group variables, we integrated the data with SIGNAL. SIGNAL successfully aligned cell types such as NK/T and B cells while distinguishing cell types like fibroblasts and smooth muscle cells across different tissues (Fig. 5 D and E and Fig. S12 A-D). Focusing on NK/T cells, which exhibit significant differences across tissues and stages, we further annotated nine subtypes based on marker genes (Fig. 5 F and G and Fig. S12E). In the SIGNAL-integrated data, T cells in the thymus were clearly distinguishable from those in other tissues, with developmental T and CD8AA subtypes being enriched in the thymus. Notably, the CD8AA subtype (marked by *GNG4*) showed differentiation across stages (Fig. 5F). We subdivided CD8AA cells into two subtypes: CD8AA I, predominantly found at 12 postconception weeks (PCW), and CD8AA II, comprising cells from multiple stages (Fig. S12F). CD8AA I overexpressed genes associated with IL-2-STAT5 signaling, such as *TNFRSF9* and *TRAF1*, while CD8AA II overexpressed genes related to TNF-α signaling via NF-κB, like *NR4A1* and *NR4A3* (Fig. S12G). We also integrated the dataset using other methods capable of incorporating additional batch variables, including Harmony, scVI, scANVI, and scPoli, with protocol as the additional technical covariate, in line with the original study. Among these methods, only scVI could preserve both tissue- and stage-specific differences, similar to SIGNAL (Fig. S12 H-K).

Lastly, we demonstrated how the SIGNAL framework can be used to construct a single-cell atlas under multiple disease conditions. We collected approximately 2.7 million human heart cells from six single-cell studies (2, 33-37), encompassing 198 donors, 229 batches, three sequencing platforms, and six disease conditions. The original data exhibited strong batch effects across different studies (Fig. S13 A-C). We integrated the data with SIGNAL using disease conditions as the group variable, performed unsupervised Leiden clustering, and annotated cell types using marker genes (Fig. 5H and Fig. S13 D and E). PGTV was then used to refine the cell labels. This workflow ultimately resulted in a well-integrated human heart cell atlas with 14 cell types, identifying several rare cell types across datasets, such as mast and epicardial cells (Fig. 5I). To test whether SIGNAL could preserve differences between disease conditions, we further examined the subtypes of fibroblasts (FB), which showed high heterogeneity under disease conditions such as dilated cardiomyopathy (DCM) and hypertrophic cardiomyopathy (HCM) in previous studies (2, 35). Using previously defined markers (33), we identified six FB subtypes, including activated FB and FB1-5 (Fig. 5 J and K). We employed Milo (38) to analyze the differential abundance of FB cell neighborhoods under disease versus healthy conditions. Our analysis revealed similar differential abundance patterns in DCM and HCM, both exhibiting an increase in activated FBs (Fig. 5K). Additionally, differentially expressed genes in activated FBs under DCM and HCM were enriched in similar pathways involved in the production, organization, and function of the extracellular matrix (ECM), as well as integrin-mediated cell adhesion and interactions (Fig. 5 K and L). Moreover, FB1 exhibited differential cell composition and distinct differential expression patterns, indicating its transcriptional heterogeneity under various disease conditions (Fig. 5K and Fig. S13F).

### SIGNAL scales to other modalities and cross-modality data

Finally, we demonstrated the scalability of SIGNAL to other modalities and cross-modality single-cell data. First, we applied SIGNAL to two single-cell proteomics datasets: the human bone marrow mononuclear cells (BMMCs) dataset (39) and the human PBMCs dataset (40) (Fig. S14 A-D). These datasets were profiled using cellular indexing of transcriptomes and epitopes by sequencing (CITE-seq) and single-cell cleavage under targets and tagmentation with cell surface proteins (scCUT&Tag-pro), respectively. Both datasets contained over 100 proteins and showed severe batch effects (Fig. S14 A and C), posing a challenging integration scenario with limited features. Our results showed that SIGNAL successfully integrated these datasets, well mixing batches and clearly separating all cell types (Fig. S14 B and D). Next, we applied SIGNAL to cross-modality data integration (Fig. S14 E-H). We utilized the human PBMCs (Multiome) dataset and the human fallopian tube (FT) dataset (41), which contained scRNA and scATAC modalities. SIGNAL successfully overcame significant modality differences, effectively integrating these cross-modality datasets. These results underscore SIGNAL’s robustness and versatility in handling diverse single-cell data types.

### Discussion

In this study, we introduced SIGNAL, a fast, robust, and flexible framework designed to interpret and remove batch effects in single-cell data by learning GTVs. This framework allows both data integration and reference-based cell label prediction. Our results demonstrated that SIGNAL performed comparably to state-of-the-art methods across a broad range of publicly available datasets.

SIGNAL integration stands out due to its simplicity and interpretability. Unlike other methods that rely on complex statistical or deep learning models, SIGNAL is a linear method requiring only basic matrix operations to project data into a batch-free low-dimensional space. Leveraging the bigstatsr R package (42), developed for facilitating large-scale matrix operations, SIGNAL can integrate large datasets of one million cells in approximately two minutes (Fig. S15).

An additional application of GTVs is the PGTV method, which allows label transfer based on reference annotations. Current workflows often involve unsupervised clustering of integrated data followed by cell type annotation based on differentially expressed genes (DEGs) or known marker genes. This process can lead to under- or over-clustering, resulting in mislabeled cells. PGTV’s self-adjustment strategy refines cell labels, restoring accurate cell grouping structures. We envisage that this strategy will be further developed to improve the clustering and annotation results in broader single-cell integrative analysis.

For reference-based integration, existing methods typically map query data to an already-integrated space before assigning labels. This can introduce unnecessary variations when the query data contains novel cell types. SIGNAL, through PGTV, assigns labels from reference data to query cells, calculates prediction confidence, and identifies potentially novel cell types. This approach allows for continuous updating of reference data and the incorporation of new biological variations. Such integration also demonstrates an ability to handle partial label input effectively. Therefore, when dealing with datasets containing millions of cells, sketching techniques can be employed to enhance computational efficiency and reduce memory load, similar to the approach in Seurat v5 (43).

SIGNAL aligns subgroups under the group variable across different batches, in contrast to methods like Harmony and scVI, which allow additional batch variables. This makes SIGNAL particularly advantageous for multi-scale analysis, especially when integrating batches with varied biological conditions. Applications of SIGNAL to three large-scale single-cell atlases confirm its suitability for customized integrative analyses.

## Materials and Methods

### SIGNAL

#### Overview

Given normalized (non-centered) single-cell data with accompanying meta information, the SIGNAL framework learns the GTVs for one or multiple group variables. Using GTVs, SIGNAL performs data integration (gcPCA) and reference-based cell labels prediction (PGTV). Below, we review the classical PCA algorithm, which forms the foundation of our framework, before detailing the SIGNAL methodology.

#### Classical PCA

Let *X* ∈ ^*M* × *N*^ denote a data matrix with *M* rows representing variables (features) and *N* columns representing observations (cells). Then *X* is centered for each variable. The centered *X* is expressed as: 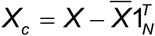, where 1_*N*_ is the *N*-dimensional vector of ones, and 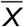 is the mean vector of *X*, i.e., 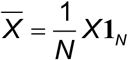. Classical PCA aims to find projection vectors *u*’s such that the total variation after projection is maximized. The optimization problem can be formulated as:

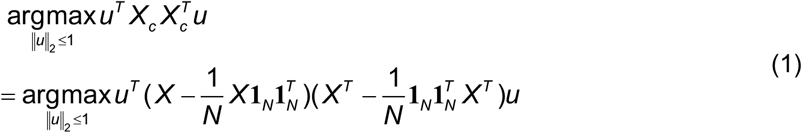

The projection vectors can be obtained as the eigenvectors of the total variatio 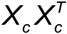.

PCA is a common approach for dimensionality reduction on a single dataset. However, it cannot deal with the batch effects present in datasets from multiple batches. Our recently proposed scRNA-seq data integration method learns the contrastive biological variation by subtracting the technical variation identified by similar cells across batches from the total variation (13). Here, we extend the concept of technical variation to one or multiple group variables, introducing GTVs.

#### GTVs

Given the data matrix *X* with a batch variable *b* (containing *L* batches) and a group variable *g* (containing *K* subgroups), let *G*_*g*_ ∈ {0,1}^*N*×*k*^ be the group-specific one-hot encoding matrix, where *G*_*g*_(*n,k*) = 1 if cell *n* is belong to subgroup *k*, and 0 otherwise. *G*_*bg*_ ∈ {0,1}^*N*×*LK*^ encodes the assignment of batches and subgroups, which is the stacked matrix of batch-group-specific one-hot encoding matrices. Let 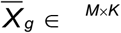 and 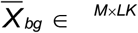 be the group means and batch-group means matrix, respectively, where column *k* of 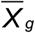 is the mean expression vector of subgroup *k* and column (*l, k*) of 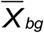 is the mean expression vector of subgroup *k* in batch *l*. We have:

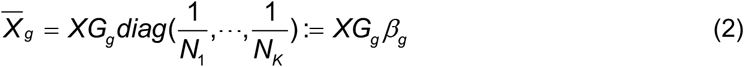

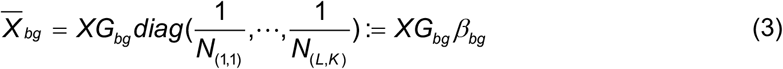

where *Nk* is the total number of cells in subgroup *k, N*(*l, k*) is the total number of cell in subgroup *k* in batch *l*, and 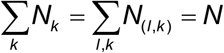. Then *X* is group-centered and batch-group-centered,

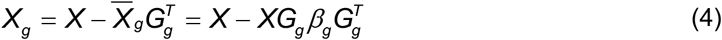

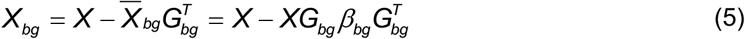

The GTV specific to the group variable *g* is:

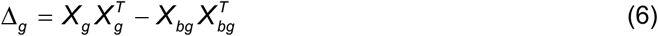

where 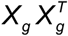 represents the sum of the overall variations under the subgroup labels, and 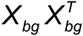 represents the sum of biological variations computed within subgroups under each individual batch label. The conceptual schematic of the composition of these variations is shown in the Fig. S1A.

#### gcPCA

We propose a novel variant of PCA, called gcPCA, which incorporates the GTVs into the PCA framework. The gcPCA model is formulated as follows:

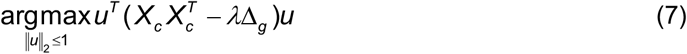

where *λ* controls the regulation strength, and *λ* = 50 by default. Similar to classical PCA, the projection vectors can be obtained as the eigenvectors of 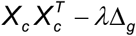.

#### gcPCA for multiple group variables

gcPCA allows multiple group variables as input. In this study, we employed two-group settings to perform reference-based integration, and unsupervised integration tasks of large-scale atlases. The decomposition of variation is illustrated in Fig. S1B. The model can be written as

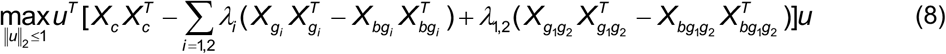

We simply set × *λ*_1_ = × *λ*_2_ = × *λ*_1,2_ (50 by default)

Given *D* group variables *g*_1_, …, *gD*, the gcPCA model can be written as

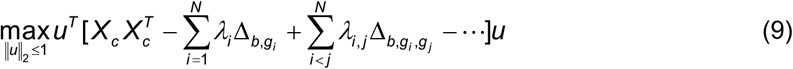

where

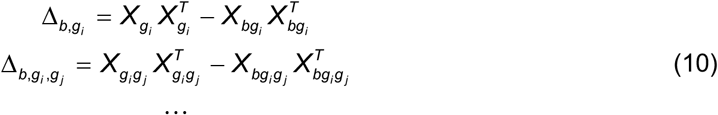

#### PGTV

The size of Δ_*g*_ is related to the subgroup label assignment of all cells in the dataset and can be formulated as:

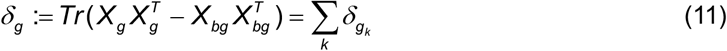

where *δ*_*gk*_ denotes the size of subgroup *k*-specific technical variation. The above equation shows that GTV can be formulated as the sum of the technical variations for each subgroup label. We assume that the technical variation is relatively small compared to the biological variation, which is a common assumption in data integration and reference-base cell type annotation. Under this assumption, given batch 1 and a single cell *i* in batch 2, if cell *i* is assigned the correct subgroup label *k*, the size of Δ_*g*_ is expected to be small. Because if cell *i* is assigned an incorrect label *k*_0_, the biological differences between cell *i* and all other cells within group label *k*_0_ in batch 1 will be perceived as part of the technical effects, resulting in a larger GTV. Thus, for cell *i*, we predict its label by finding the subgroup label *k* that minimizes the incremental technical variation of *k*:

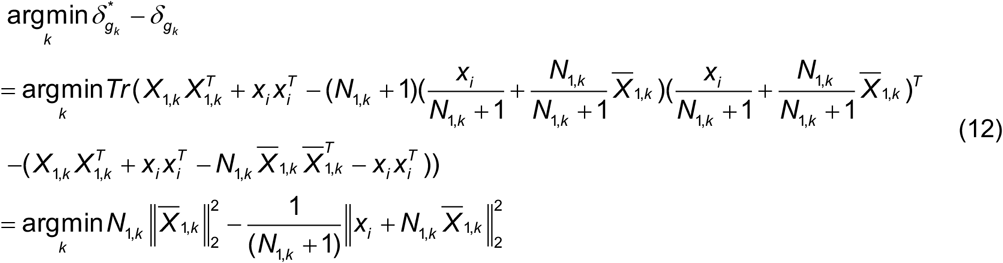

where *N*_1,*k*_, 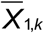 represent the cell number and mean expression vector of subgroup *k* of batch 1, respectively, and *xi* is the expression vector of cell *i* in batch 2. So far, we formulated the label prediction problem to a simple classification problem.

#### Confidence score of label prediction

Above cell label prediction procedure implicitly assumes that cell *i* belongs to a cell type present in batch 1. However, it is crucial to account for the situation that cell *i* represents a novel cell type, which is common in single-cell data analysis. Thus, we introduce a confidence score of reference-based cell label prediction. After cell label prediction from reference to query, we use the cosine similarities between reference group center vectors and query cells in PCA space as proxies of the confidence scores. Specifically, for a reference dataset *X*_*ref*_ and a query dataset *X*_*query*_, we compute the gene means *μ* and standard deviations *δ* of *X*_*query*_. These statistics are used to center and scale both *X*_*ref*_ and *X*_*query*_. Using singular value decomposition (SVD), we learn the first *d* left singular vectors of *X*_*query*_, enabling the projection of *X*_*query*_ and centroids of *K* groups in *X*_*ref*_ into the same PCA space. Then, we calculate the cosine similarities *Ci* in the low-dimensional space between each cell *i* in *X*_*query*_ and the centroid of its assigned group *k*. To smooth the confidence score, *Ci* is computed as the average confidence score of its *smooth*.*k* (20 by default) nearest neighbor query cells. We introduce a confidence cell ratio,*η* (0.2 by default), representing the proportion of high-confidence cells in the query dataset.

The confidence scores are then scaled between 0 and 1 using the 0.01 and 1 − *η* percentiles.

#### Cell label self-adjustment strategy

Given a dataset and its cell labels *L*_0_, we use PGTV to obtain predicted labels *L*_1_ based on *L*_0_. Then, we calculate the adjusted Rand index (ARI) value or any other measure of label consistency, *a*_1_, between *L*_0_ and *L*_1_. If *a*_1_ < 1, we predict the cell labels using the overlapping labels between *L*_0_ and *L*_1_ to obtain adjusted labels *L*_2_, and calculate the ARI value *a*_2_ between *L*_1_ and *L*_2_. This step will be iteratively repeated until the difference between *at* and *at-1* (where *t* is the iteration number) is less than a predefined threshold (0.001 by default).

#### Reference-based integration

For an annotated reference dataset and an unannotated query dataset, PGTV is used to predict the cell labels and calculate confidence scores for the query data. Then, cell labels from the query with confidence scores greater than *μ* = 0.8 and all cell labels from the reference are used as inputs for gcPCA. Additionally, we introduce a new group variable, ‘dataset’, which includes two subgroups: ‘reference’ and ‘query’, representing cells belonging to the reference and query datasets, respectively. The cell label and the dataset group variables were used to perform gcPCA.

#### SIGNAL parameter

The SIGNAL framework has a single tuning parameter, *λ*, which was set to 50 for all integrative analyses in this study. We tested SIGNAL integration results with different *λ* values on the human pancreas dataset and found that SIGNAL exhibited robust performance across various *λ* choices (Fig. S16).

#### Runtime and memory scalability analysis

We subsampled the HLCA dataset proposed by Sikkema et al. (32), and got six datasets with cell numbers ranging from 20,000 to one million with 2,000 highly variable genes (HVGs). The runtime and peak RAM usage of the SIGNAL integration were measured using the peakRAM R package (v1.0.2). All analyses were conducted on a Linux machine equipped with a 1.50GHz AMD EPYC 7513 processor and 256 GB of memory.

### Datasets

#### Data integration benchmark scRNA-seq datasets

We utilized eight scRNA-seq datasets from two recently published benchmark studies (4, 14), comprising six real datasets (human DCs, cell lines, human pancreas, human lung, human immune, and immune (cross-species) datasets) and two simulated datasets (Sim 1 and Sim 2). Processed data were obtained from the respective study websites. Additionally, we collected the WAT (cross-species) dataset from a recent cross-species study (15), which includes 36 batches and 363,870 cells from human and mouse white adipose tissue. The biomaRt R package (v2.54.1) was employed to map mouse genes to human genes, resulting in the retention of 15,530 shared genes across species.

#### Label prediction benchmark scRNA-seq datasets

We assessed cell type prediction performance using two simulated datasets (Sim 1 and Sim 2) and four real datasets (human cortex, HLCA_Travaglini, mouse retina, and Tabula Muris atlas) (see Table S1). Specifically, for human cortical data, we collected two human cortical datasets from different studies. The first dataset consists of snRNA-seq data from the human MTG, containing five batches and 137,303 nuclei. The second dataset contains 379,330 nuclei across 78 batches from the human cortex. We utilized the human MTG dataset to predict cell type labels for the human cortex dataset. For the HLCA_Travaglini dataset, we used three batches (containing 65,662 cells) assayed with 10x Genomics to predict the cell type labels of three batches (containing 9,268 cells) from Smart-seq2 (SS2). For the mouse retina dataset, we used the ‘Shekhar’ batch (containing 26,830 cells) to predict the other batch (containing 43,603 cells). For the Tabula Muris atlas, we used cells from the 3-month time point (containing 90,120 cells) to predict all other cells (containing 266,093 cells). For all real datasets, the cell type annotations from the original studies were used as the ground-truth cell type labels. For the simulated datasets, we simply used half of the batch to predict the cell labels for the other half.

#### Single-cell proteomics datasets

We curated the proteomics data from the CITE-seq dataset included in the NeurIPS 2021 multimodal data integration datasets (39) to construct the human BMMCs dataset. For the human PBMCs dataset, we used the proteomics data from six scCUT&Tag-pro datasets as provided in the original study (40).

#### Cross-modality single-cell datasets

The human PBMCs dataset was obtained from the SeuratData R package (v0.2.2) and includes scRNA and scATAC modalities measured in the same cells. We treated these two modalities as if they originated from separate batches to test SIGNAL’s capability in integrating cross-modality data. For the human FT dataset, we utilized the normalized gene activity matrix for scATAC-seq data provided by the original study (41). For both datasets, we selected HVGs in the scRNA-seq batches as the features for integration.

### Analysis details

#### Data pre-processing

For the human heart data generated by Reichart et al. (35) and the HLCA dataset proposed by Sikkema et al. (32), we removed cells labeled as “unassigned”. For single-cell datasets curated by Luecken et al. (4), we directly used the normalized data provided by the authors. For other scRNA-seq datasets, we used the count data provided by the original studies and performed standard normalization procedures using the Seurat R package (v4.3.0.1). Except for the developing human immune dataset, where we used the HVGs set provided by the authors, we selected HVGs using the FindVariableFeatures function in Seurat. For cross-species datasets, HVGs were selected separately for each species and then merged. For the human pancreas dataset, which came from multiple sequencing technologies, we scaled the data to reduce the impact of sequencing platform differences. For all scRNA-seq datasets, we used their HVGs for subsequent analyses.

#### Data visualization

UMAP embeddings for all datasets were generated using the same parameters using the uwot R package (v0.2.2), with *min_dist* = 0.25, except for the human heart cell atlas dataset, which *min_dist* = 0.1 was used.

#### Benchmark data integration methods

We benchmarked SIGNAL integration against nine other data integration methods: fastMNN, Seurat, Harmony, LIGER, BBKNN, Conos, scVI, scANVI, and scPoli. For all methods except LIGER, we set a unified dimensionality *d* for the integrated low-dimensional space within the same dataset, typically 30 or 50, with *d* set to 50 for large datasets. For LIGER, the default *k* value of 20 was used. For Seurat, the CCA mode was employed unless it could not be executed due to large datasets, in which case the RPCA mode was used. Apart from these considerations, all methods were run with their default parameters for integration. These methods were performed using batchelor (v1.14.1 for fastMNN), Seurat (v4.3.0.1), harmony (v0.1.1), rliger (v1.0.0 for LIGER), and conos (v1.5.1) in R, and bbknn (v1.6.0), scvi-tools (v1.0.4 for scVI and scANVI), and scArches (v0.5.9 for scPoli) in Python.

#### Benchmark reference-based annotation methods

We benchmarked PGTV against four reference-based annotation methods: SingleR, SciBet, scmap, and scPred. For scmap, both clustering-based and nearest-neighbor-based strategies (scmap-cluster and scmap-cell) were employed. All methods used the reference dataset with its ground truth labels as input, and default parameters were employed to predict cell type labels for the query dataset. These methods were performed using SingleR (v2.0.0), scibet (v1.0), scmap (v1.20.2), and scPred (v1.9.2) in R.

#### Benchmark reference-based integration methods

We benchmarked SIGNAL against four reference-based integration methods: scANVI, scPoli, Seurat and Symphony. All methods were applied using default parameters for reference-based integration. These methods were performed using scvi-tools (v1.0.4) and scArches (v0.5.9) in Python, and Seurat (v4.3.0.1) and symphony (v0.1.1) in R.

#### Evaluation metrics

To assess integration performance, we employed evaluation metrics summarized by Luecken et al. (4). These metrics, calculated using the scib-metrics Python package (v0.4.1), include measures of biological variation conservation (isolated label score, normalized mutual information (NMI), ARI, average silhouette width (ASW) label, and cLISI) and batch effect removal (ASW batch, iLISI, kBET, and graph connectivity). Additionally, batch correction scores, biological variation conservation scores, and overall integration scores were calculated. To ensure fair comparisons, as graph-based methods do not produce low-dimensional matrices, we used UMAP embeddings as inputs. For reference-based annotation tasks, performance was evaluated using common metrics including accuracy and macro F1. In our previous work, we used accuracy and F1 scores to assess the cell label prediction performance of reference-based integration (13). The macro F1 score, being the average F1 score across all cell types, is more sensitive to rare cell types. In reference-based integration tasks, alongside accuracy and macro F1 for assessing label transfer accuracy, we also used our previously proposed modified LISI to measure mixing across different datasets (dataset LISI) and developmental stages (age LISI). Details for all these metrics can be found in ref. 4 and ref. 13.

#### K-means clustering and supervised integration of the human pancreas dataset

We performed k-means clustering on both the unintegrated and unsupervised integrated data, with *k* set to 14. The clustering results were then used as inputs for the supervised integration methods. ARI values between the clustering results and the ground truth cell type labels were also calculated.

#### Label self-adjustment analysis

To assess the proposed label self-adjustment strategy, we simulated mislabeled scenarios on the Sim 1 dataset, where ground truth labels were known. For each batch in Sim 1, we identified the cell type *i* with the largest number of cells and its nearest neighbor cell type *j*, and artificially mislabeled 50% of the cells closest to *j* in *i*. This generated a mislabeled dataset. We applied PGTV label self-adjustment separately on individual batches and on the entire dataset. For the human PBMCs dataset, we obtained new labels using PGTV label self-adjustment. We identified the set of cells *A* with inconsistent labels between the original labels and the predicted labels, and the set of cells *B* with consistent labels. For each cell in set *A*, we calculated the cosine similarity between its gene expression and the mean gene expression assigned by different methods in set *B*. Additionally, we compared the average log-fold change of the top five cell type markers between the original labels and the labels adjusted by PGTV.

#### Developing human immune cell atlas analysis

For the developing human immune cell atlas dataset, we utilized 3,765 HVGs provided by the original study. The cell type labels provided with the data were used for scANVI and scPoli integration.

#### Human heart cell atlas analysis

We compiled human cardiac datasets from six studies, containing a total of 2,747,449 cells. For each study, we computed the top 2,000 HVGs individually and then selected a final set of 2,000 HVGs for analysis using the SelectIntegrationFeatures function in the Seurat. Donor and condition were used as the batch and group variables, respectively, for SIGNAL integration, resulting in a 50-dimensional gcPCA embedding. The entire integration process took approximately 10 minutes.

#### Statistical analysis

Differential expression analysis was performed using the Wilcoxon rank-sum test, implemented through the FindMarkers function in Seurat. GO enrichment analysis was conducted using the clusterProfiler R package (v4.6.2) (44), with *P* value and *Q* value thresholds set at 0.05, and *P* values adjusted using the Benjamini-Hochberg method.

## Data availability

All datasets used in this manuscript are publicly available. Details of the datasets are listed in Table S1.

## Code availability

The SIGNAL R package is available freely at GitHub (https://github.com/yzhou1999/SIGNAL). Source codes to reproduce the results in this manuscript are available at GitHub (https://github.com/yzhou1999/SIGNAL_reproducibility).

## Acknowledgments

This work was supported by the National Natural Science Foundation of China (Grants No. 62271173 and No. 62172122), the Key Research and Development Program of Heilongjiang (Grant No. 2022ZX01A19), and the Natural Science Foundation of Heilongjiang Province, China (Grant No. JQ2023A003). We thank Servier Medical Art (https://smart.servier.com) for the schematics.

